# A distinct Down state assembly in retrosplenial cortex during slow-wave sleep

**DOI:** 10.1101/2024.07.19.604325

**Authors:** Ashley N Opalka, Kimberly J Dougherty, Dong V Wang

## Abstract

Understanding the intricate mechanisms underlying slow-wave sleep (SWS) is crucial for deciphering the brain’s role in memory consolidation and cognitive functions. It is well-established that cortical delta oscillations (0.5–4 Hz) coordinate communications among various cortical, hippocampal, and thalamic regions during SWS. These delta oscillations have periods of Up and Down states, with the latter previously thought to represent complete cortical silence; however, new evidence suggests that Down states serve important functions for information exchange during memory consolidation. The retrosplenial cortex (RSC) stands out for its pivotal role in memory consolidation due to its extensive connectivity with memory-associated regions, although it remains unclear how RSC neurons engage in delta-associated consolidation processes. Here, we employed multi-channel *in vivo* electrophysiology to study RSC neuronal activity in freely behaving mice during natural SWS. We discovered that the RSC contains a discrete assembly of putative excitatory neurons (∼20%) that initiated firing at SWS Down states and reached maximal firing at the Down-to-Up transitions. Therefore, we termed these RSC neurons the Down state assembly (DSA), and the remaining RSC excitatory neurons as non-DSA. Compared to non-DSA, DSA neurons exhibit a higher firing rate, larger cell body size, and no connectivity with nearby RSC neurons. Subsequently, we investigated RSC neuronal activity during a contextual fear conditioning paradigm and found that both DSA and non-DSA neurons exhibited increased firing activity during post-training sleep compared to pre-training sleep, indicating their roles in memory consolidation. Lastly, optogenetics combined with electrophysiology revealed that memory-associated inputs differentially innervated RSC excitatory neurons. Collectively, these findings provide insight on distinct RSC neuronal subpopulation activity in sleep and memory consolidation.

## INTRODUCTION

Sleep is a vital physiological process that constitutes about one third of human life. A significant portion of sleep, known as slow-wave sleep (SWS), is characterized by delta oscillations (0.5–4 Hz), featuring cyclic fluctuations between active periods and silent periods, known as Up and Down states, respectively. This rhythmic activity coordinates various brain regions to optimize communication, playing a pivotal role in memory consolidation, a process that converts hippocampal-dependent memories into long-term cortex-dependent memories [1-8]. In support of this, studies have demonstrated that boosting the cortical delta oscillation during post-learning sleep enhanced memory retention [4, 9]. Additionally, strengthening interactions between delta and higher-frequency oscillations, such as hippocampal sharp-wave ripples or thalamocortical spindles, also improves memory [7, 8, 10, 11].

The precise mechanism through which delta oscillations initiate information flow and facilitate memory consolidation has yet to be fully elucidated. Recent work indicates that memory consolidation involves a cortical-hippocampal-cortical loop of information processing, highlighting the important role of the cortex in initiating the consolidation process [6, 12-14]. It is plausible that Up and Down states serve distinct functions, with Down states acting as a reset mechanism to prepare key brain regions for efficient information exchange during subsequent Up states, thus facilitating the consolidation of memory episodes.

The retrosplenial cortex (RSC) emerges as a key cortical region for memory consolidation, given its dense connections with various memory-associated regions, including the hippocampal formation, anterior thalamus, and other cortical regions [15-21]. Studies have shown that disruption of the RSC immediately after learning impairs memory, while activation of the RSC during post-learning sleep enhances memory [14, 22-24]. Furthermore, recent findings underscore the significance of hippocampus-RSC interactions in memory consolidation, highlighting the RSC’s role as a link to integrate hippocampal, cortical, and subcortical regions to facilitate memory processing [18-20]. However, the specific involvement of distinct RSC neuronal populations during memory consolidation remains unclear.

Here, we aimed to identify patterns of RSC activity during SWS and discovered a distinct subpopulation of RSC neurons that are active during SWS Down states and Down-to-Up transitions. We further explored how RSC neurons participate in SWS-dependent memory consolidation and receive innervation from distinct memory-associated inputs. Our findings underscore the RSC as a key structure in orchestrating memory consolidation.

## RESULTS

### Characterizing a novel RSC Down State Assembly (DSA) during SWS

To investigate RSC neuronal and oscillatory activity, we performed extracellular recordings mainly in RSC deep layers of freely behaving mice (8–16 tetrodes) [18, 25]. Our objective was to first organize RSC population activity into neuronal assemblies, subgroups of RSC neurons that exhibited co-activation. To achieve this, we employed a combined principal and independent component analysis (PCA–ICA) [26], focusing on natural slow-wave sleep (SWS) [18], where the RSC displays cyclic Up-Down patterns of activity (Figure 1A&B; Suppl. figure 1A&B). Of the ICA-detected RSC assemblies across 8 animals (n = 36; 3.6 ± 1.1 assemblies, mean ± SD; ranged between 2 and 6 per recording session), we noticed that one distinct assembly of RSC neurons from each recording session exhibited high co-activity during SWS Down states, as well as Down-to-Up transitions (Figure 1A). We thus termed this assembly the RSC Down State Assembly (DSA).

**Figure 1:**
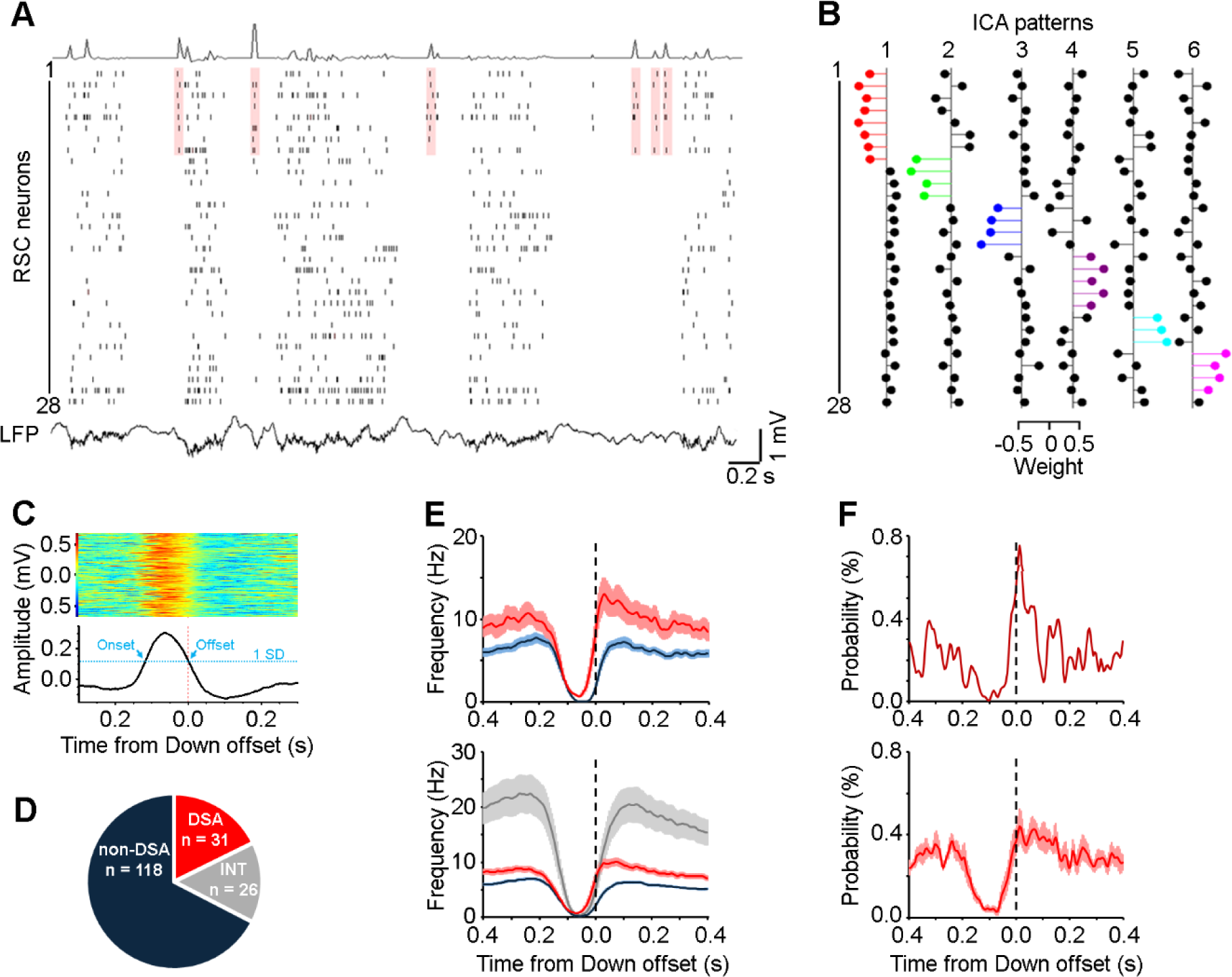
RSC DSA activity precedes local neuronal activity at Down-to-Up transition. (A) Raster plot of 28 neurons and local field potential (LFP) recorded simultaneously from RSC during SWS. Shading highlights DSA activity (neurons #1–8) during Down states when non-DSA neurons (#9–26) and putative interneurons (#27, #28) are mostly silent. Top, DSA activity strength estimated by principal components. (B) An example independent component analysis (ICA) identified six assembly patterns based on activity of 28 RSC neurons organized in the same sequence as A. (C) Representative Down states (top) and averaged (bottom) in reference to Down offset, which indicates the timepoint of mean + SD at the descending phase. (D) Percentage of DSA excitatory, non-DSA excitatory, and interneurons. (E) Mean frequency (± SEM) of DSA neurons (red) precedes non-DSA neurons (navy) and interneurons (grey) in reference to RSC Down offset from one mouse (top) and across mice (bottom; n = 10). (F) Cross-correlograms between Down offsets and DSA activity from one mouse (top) and across mice (bottom; n = 10; mean ± SEM). E&F bin size = 5 ms. All examples were simultaneously recorded from the same mouse.

To further characterize how RSC neurons fire in relation to Down states, we calculated Down state offsets (and onsets) as when the RSC delta oscillation (bandpass filtered between 1–4 Hz) amplitudes surpassed one standard deviation (SD) above the mean (Figure 1C; see Methods for details). Additionally, we classified our recorded RSC neurons into either putative excitatory neurons or putative interneurons based on spike waveform using an unsupervised principal component analysis (PCA; 149 and 26 out of 175, respectively; Suppl. figure 2) [18]. Our results suggest that all DSA neurons are putative excitatory neurons. Accordingly, the remaining RSC putative excitatory neurons will be referred to as non-DSA neurons.

Our correlation analysis revealed that RSC DSA neurons initiated activity during Down states and reached maximal activity at Down-to-Up transitions, which preceded the activity of RSC non-DSA neurons (Figure 1E, top). On average, DSA mean firing activity (red; 31/175; maximum firing of 10.12 Hz at 0.07 s) not only preceded the activity of non-DSA neurons (navy; 118/175; maximum firing of 6.36 Hz at 0.10 s) but also interneurons (grey; 26/175; maximum firing of 20.4 Hz at 0.12 s) across mice (Figure 1E, bottom). Notably, this DSA activity was not observed in other midline cortices, including the anterior cingulate cortex and prelimbic cortex (Suppl. figure 3A). Consistently, DSA co-activity strength, calculated based on by principal components [26] (as seen in Figure 1A, top), exhibited the highest probability of co-activity at the Down-to-Up transition (0.01 s for both the example and average across mice; Figure 1F). These findings highlight the distinctness of RSC DSA activity at SWS Down-to-Up transitions.

Overall, the DSA represents about 21% of the recorded neuronal population (20.81 ± 3.15%, mean ± SD) and was observed in 80% of mice (8/10), largely in the deep-layer (layer 5) granular RSC (Suppl. figure 1C&D). In addition, the DSA was robustly and reliably detected by the PCA–ICA, regardless of whether RSC interneurons were included for the analysis or the selection of co-firing time windows (10, 25, or 40 ms; Suppl. figure 4A). To our knowledge, this is the first identification of an assembly of cortical neurons that co-fire during SWS Down states.

### DSA neurons exhibit distinct firing, morphology, and connectivity properties

To further characterize DSA activity during SWS Down states, we computed the probability of RSC neurons across Down states. Our results revealed that DSA neurons had significantly higher firing probability during Down states compared to non-DSA neurons or interneurons (*H*_(2)_ = 30.702, p < 10^-7^, Kruskal–Wallis test; Figure 2A), along with other cortical neurons (Suppl. figure 3B). Additionally, DSA neurons exhibited significantly higher firing rates during SWS compared to non-DSA neurons (*H*_(2)_ = 42.748, *p* < 10^-10^, Kruskal–Wallis test; Figure 2B). Overall, these findings indicate that DSA neurons are more active during SWS, particularly during SWS Down states that are typically silent.

**Figure 2:**
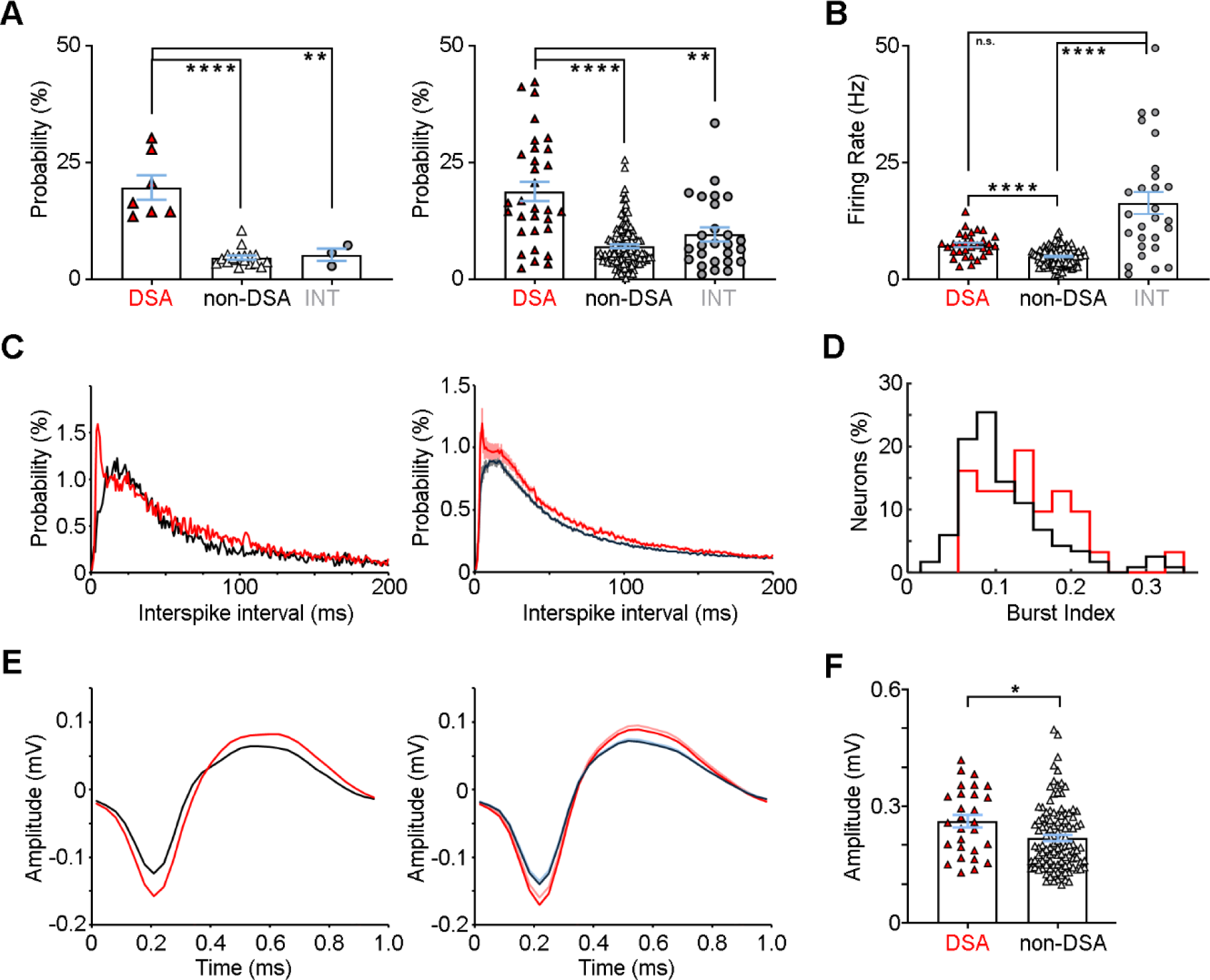
DSA neurons are a distinct subset of excitatory neurons. (A) DSA neurons exhibit significantly greater down state firing probability compared to non-DSA neurons and interneurons from one mouse (left) and across mice (right; n = 10). (B) DSA neurons have significantly greater firing rates than non-DSA neurons. (C) Inter-spike interval of DSA neurons (red) and non-DSA (black) neurons from one mouse (left, examples) and across mice (right; n = 10; mean ± SEM). (D) DSA neurons have significantly higher burst indices compared to non-DSA neurons. (E) Spike waveforms of DSA (red) and non-DSA (black) neurons from one mouse (left, examples) and across mice (right; n = 10; mean ± SEM). (F) DSA neurons have significantly greater spike amplitude compared to non-DSA neurons. All examples were simultaneously recorded from the same mouse. Error bars = mean ± SEM.

Our analysis also revealed that DSA neurons exhibited different bursting properties. Specifically, the peak inter-spike interval (ISI) of DSA neurons was significantly shorter compared to non-DSA neurons (z = -2.677, *p* < 0.01; Wilcoxon rank sum test; Figure 2C; Suppl. figure 3C). We further calculated the burst index defined as the percent of spikes with ISI < 10 ms (see Methods) and found that DSA neurons had significantly greater burst indexes compared to non-DSA neurons (z = -2.670, *p* < 0.01; Wilcoxon rank sum test; Figure 2D). Furthermore, DSA neurons displayed significantly larger spike amplitudes compared to non-DSA neurons (z = -2.460, p = 0.014, Wilcoxon rank sum test; Figure 2E&F; Suppl. figure 3D; see Methods), suggesting larger cell body sizes of DSA neurons [27]. These collective properties indicate that the DSA consists of a unique subpopulation of RSC neurons.

Next, we investigated the local connections amongst RSC subpopulations. Cross-correlation analysis revealed a reciprocal monosynaptic connection between non-DSA (excitatory) and interneurons (Figure 3A), indicated by an inhibition (between –2 and –1 ms) followed by activation (between 1 and 2 ms) or vice versa (Suppl. figure 5). This is consistent with previous work in the RSC [28, 29].

**Figure 3:**
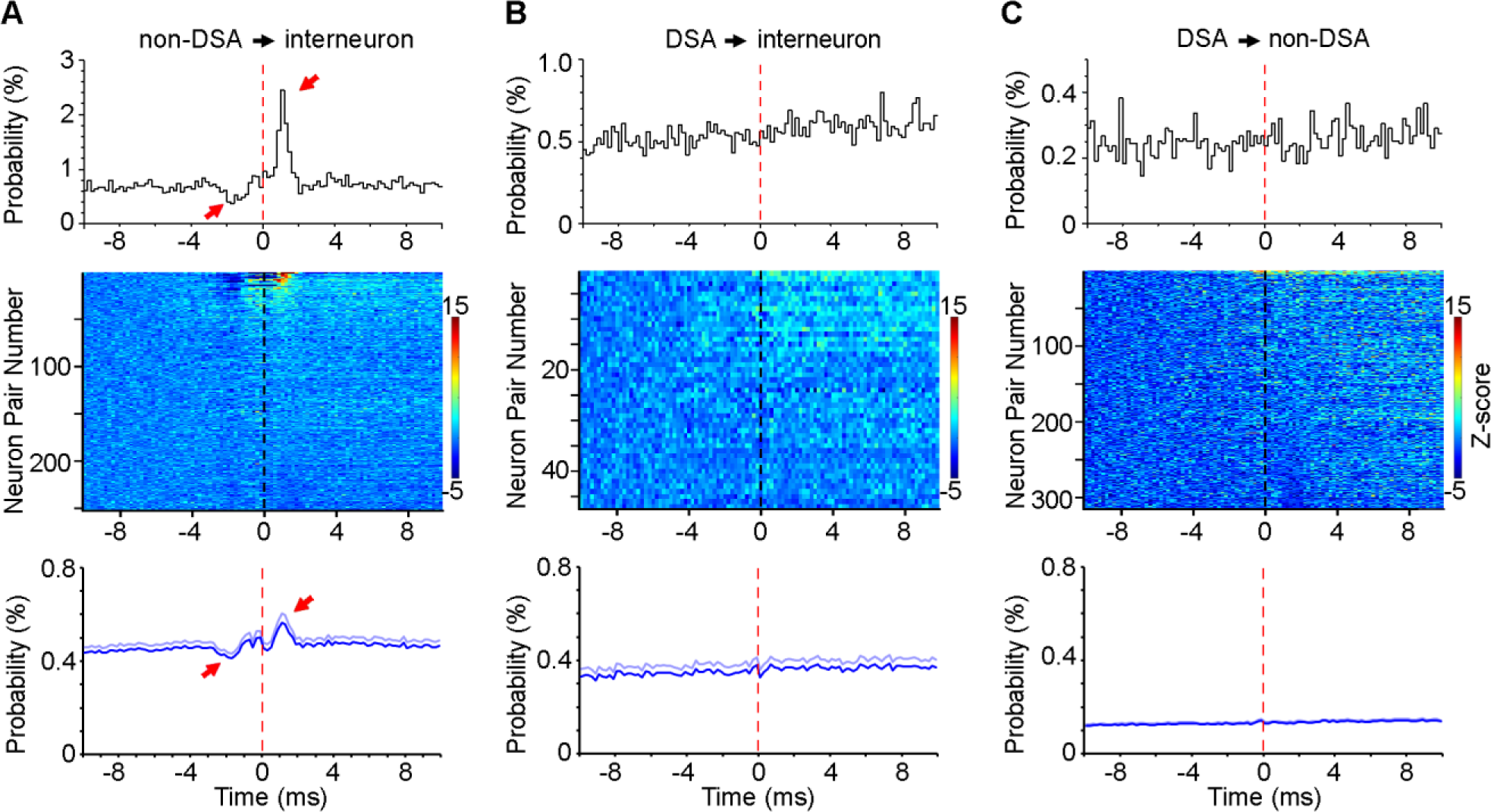
DSA neurons have weak local interactions. (A–C) Cross-correlations between RSC neuronal subpopulations. Neuron pairs include non-DSA neurons and interneurons (A), DSA neurons and interneurons (B), and DSA and non-DSA neurons (C). Interneurons were used as the reference in A and B, while non-DSA neurons were the reference in C. Top panels, representative neuronal pairs. Middle panels, heatmap of individual neuron pairs. Color bar indicates Z-scored firing probability. Neuron pairs are arranged from high-to-low firing between 1–2 ms, and Z-score transform was calculated between -10 and -5 ms based on mean and SD. Bottom panels, mean ± SEM of neuronal pairs. Only neuron pairs recorded from different tetrodes were used in this analysis.

In contrast, our cross-correlation analyses revealed that DSA neurons displayed a weak, if any, connection with interneurons (Figure 3B). Meanwhile, cross-correlations between DSA and non-DSA neurons revealed no monosynaptic connection (Figure 3C). This suggests that DSA neurons do not directly communicate with other RSC deep-layer neurons; instead, they are likely connected to RSC neurons in superficial layers and/or extra-RSC brain regions.

### DSA and non-DSA neurons exhibit activity and pattern changes during memory consolidation

In a subset of mice (n = 6), we conducted contextual fear conditioning (CFC) to ascertain the role of RSC neurons in learning and memory processes. Specifically, we focused on post-training SWS, a crucial time window for memory consolidation [30, 31]. We recorded RSC neuronal activity of mice during pre-training sleep, CFC training, post-training sleep, and contextual recall (re-exposure to the footshock chamber; Figure 4A). We focused our analyses on DSA and non-DSA neurons due to the limited number of interneurons (n = 8).

**Figure 4:**
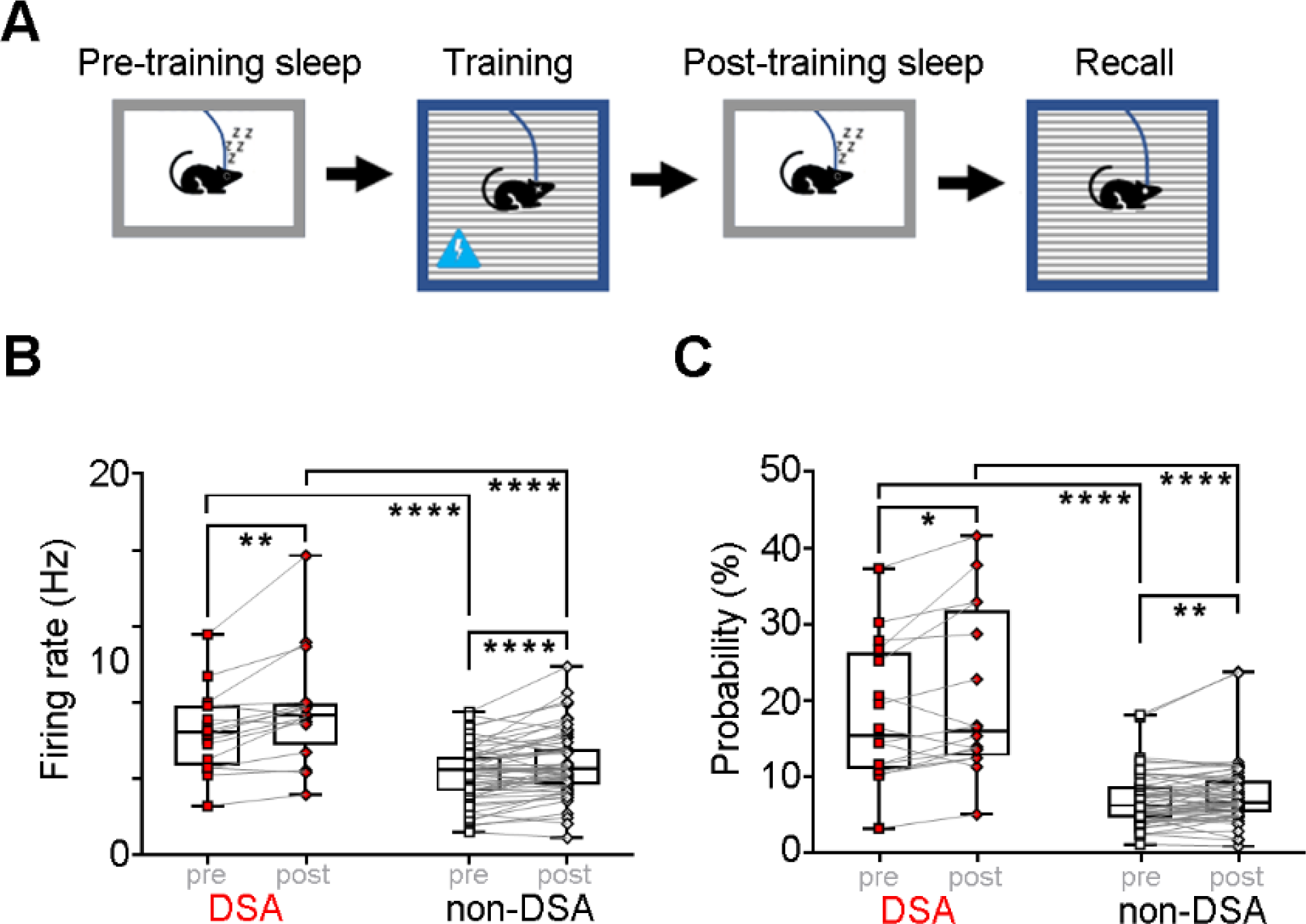
DSA and non-DSA neurons in contextual fear conditioning. (A) Schematic of contextual fear conditioning paradigm. (B&C) DSA and non-DSA neurons have significantly different firing rates (B) and down state firing probability (C) from each other and between pre- and post-training SWS sessions.

We found that DSA and non-DSA neurons had significantly greater firing rates between pre- and post-training SWS (F_(1, 69)_ = 36.382, p < 10^-8^, mixed ANOVA; Figure 4B), indicating their important roles in memory consolidation. Additionally, there is a significant difference of firing rates between DSA and non-DSA neurons (F_(1, 69)_ = 24.521, p < 10^-6^, mixed ANOVA), consistent with earlier results shown in Figure 2. We found a similar phenomenon when investigating Down state firing probability, which was significantly greater during post-training sleep compared to pre-training sleep for both DSA and non-DSA neurons, DSA: z = -2.017, p < 0.05; non-DSA: z = 2.807, p < 0.01, Wilcoxon signed-rank test; Figure 4C) and significantly greater in DSA neurons compared to non-DSA neurons at each time point (DSA: z = -4.790, p <10^-6^; non-DSA: z = - 5.120, p < 10^-7^; Wilcoxon rank sum test).

Furthermore, we found that the membership of DSA neurons showed little change across sleep sessions, regardless of the animal’s history of receiving CFC training (Suppl. figure 4B). This suggests that the RSC DSA is a preconfigured (or hard-wired) assembly of neurons, and its activity may provide structural support (i.e., promoting SWS Up states) rather than encoding specific information.

Collectively, these results reveal that both RSC DSA and non-DSA neurons display significantly greater firing activity and Down-state firing probability during post-training sleep, a crucial window in memory consolidation.

### Optostimulation of RSC inputs differentially influence RSC excitatory neurons

DSA neurons fire in high synchrony during SWS Down states and Down-to-Up transitions, raising the possibility of a shared input to these DSA neurons associated with Down states. Two possible candidates that emerge are the anteroventral thalamus (AVT) and claustrum (CLA), both of which project directly to the RSC and engage in the generation of cortical delta oscillations [3, 32]. To this end, we utilized a combined optogenetic and electrophysiological approach: we injected retrograde AAV viruses into the RSC of mice and then implanted a tetrode array in the granular RSC and an optic fiber in either the AVT or CLA ipsilaterally (Figure 5A&B). This allows us to investigate the optostimulation of the AVT-to-RSC and CLA-to-RSC pathways on RSC activity. For comparisons, we included optostimulation of the dHPC-to-RSC pathway here from our previous work [18].

**Figure 5:**
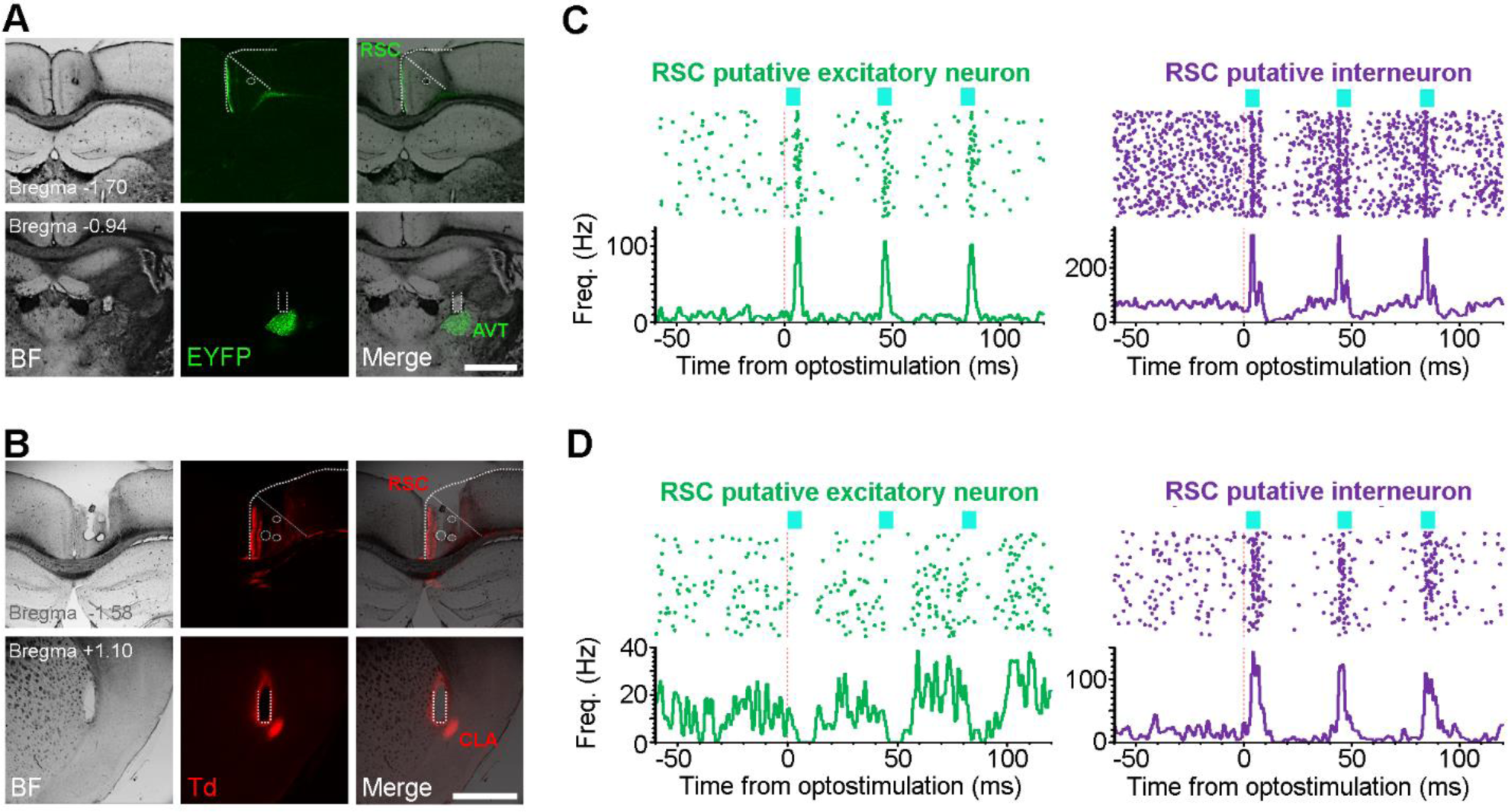
AVT and CLA regulate RSC neuronal firing. (A&B) Top, Representative coronal sections showing final recording sites (dashed circles) and viral expression in granular RSC. Bottom, retrograde viral labeling of reporter protein in AVT (A) and CLA (B) and optic fiber tracks (dashed lines). Scale bars, 1 mm. (C&D) Representative peri-event rasters of AVT (C) and CLA (D) cell body optostimulation that activated and suppressed RSC putative excitatory neurons (C&D, left, respectively) and activated interneurons (right). RSC neurons shown in C were recorded simultaneously; neurons shown in D were also recorded simultaneously. Blue bars: three laser pulses at 25 Hz; pulse width, 3 ms; n = 120 trails in C, n = 124 trials in D.

When investigating the AVT-to-RSC pathway, we opted to use Shox2-Cre transgenic mice to selectively target the thalamocortical pathway while excluding the corticothalamic pathway, given that Shox2 expression is restricted to cell bodies of the thalamus but not cortex [33]. First, we characterized the thalamic Shox2 input to cortex via the injection of Cre-On/-Off retrograde AAVrg-Ef1a-DO_DIO-TdTomato_EGFP-WPRE-pA [34] into the RSC of adult Shox2-Cre mice [35]. This Cre-On/Off AAVrg enables the expression of EGFP when Cre is present and TdTomato when Cre is absent. Our results revealed EGFP+ cell bodies densely located in the AVT and sparsely in the neighboring anterodorsal and anteromedial thalamus (ADT/AMT; Figure 5A; Suppl. figure 6&7). Remarkably, none of the AVT or ADT/AMT neurons expressed TdTomato, indicating that all AVT^➔RSC^ neurons express Shox2. This finding was further verified by inspection of another major RSC input, the subiculum, which expressed dense TdTomato but no EGFP (Suppl. figure 7). Therefore, the use of the Shox2-Cre line enables precise targeting of the AVT-to-RSC pathway, while simultaneously recording RSC neuronal activity (Figure 5C).

To investigate how the CLA influences RSC activity, we used a similar retrograde targeting strategy in wildtype mice, which enabled specific targeting of the CLA-to-RSC pathway while simultaneously recording RSC neuronal activity (Figure 5B&D). Notably, optogenetic stimulation was selective to the CLA, as surrounding regions do not project to the RSC.

We first investigated how RSC putative excitatory neurons are influenced by optostimulation of these pathways (Figure 6). Upon optostimulation of the AVT-to-RSC pathway, one third of RSC putative excitatory neurons (33.3%, 23/69; Figure 6A) were significantly activated. The response latency – defined as latency to peak firing rate – was short for most RSC putative excitatory neurons (6.4 ± 1.5 ms, mean ± SD; n = 18), indicating a monosynaptic projection from AVT to RSC excitatory neurons. Meanwhile, a small portion of RSC putative excitatory neurons (n = 5) had a longer response latency (13.7 ± 3.4 ms, mean ± SD; n = 5; ranged between 11 and 20 ms), likely indicative of a polysynaptic response.

**Figure 6:**
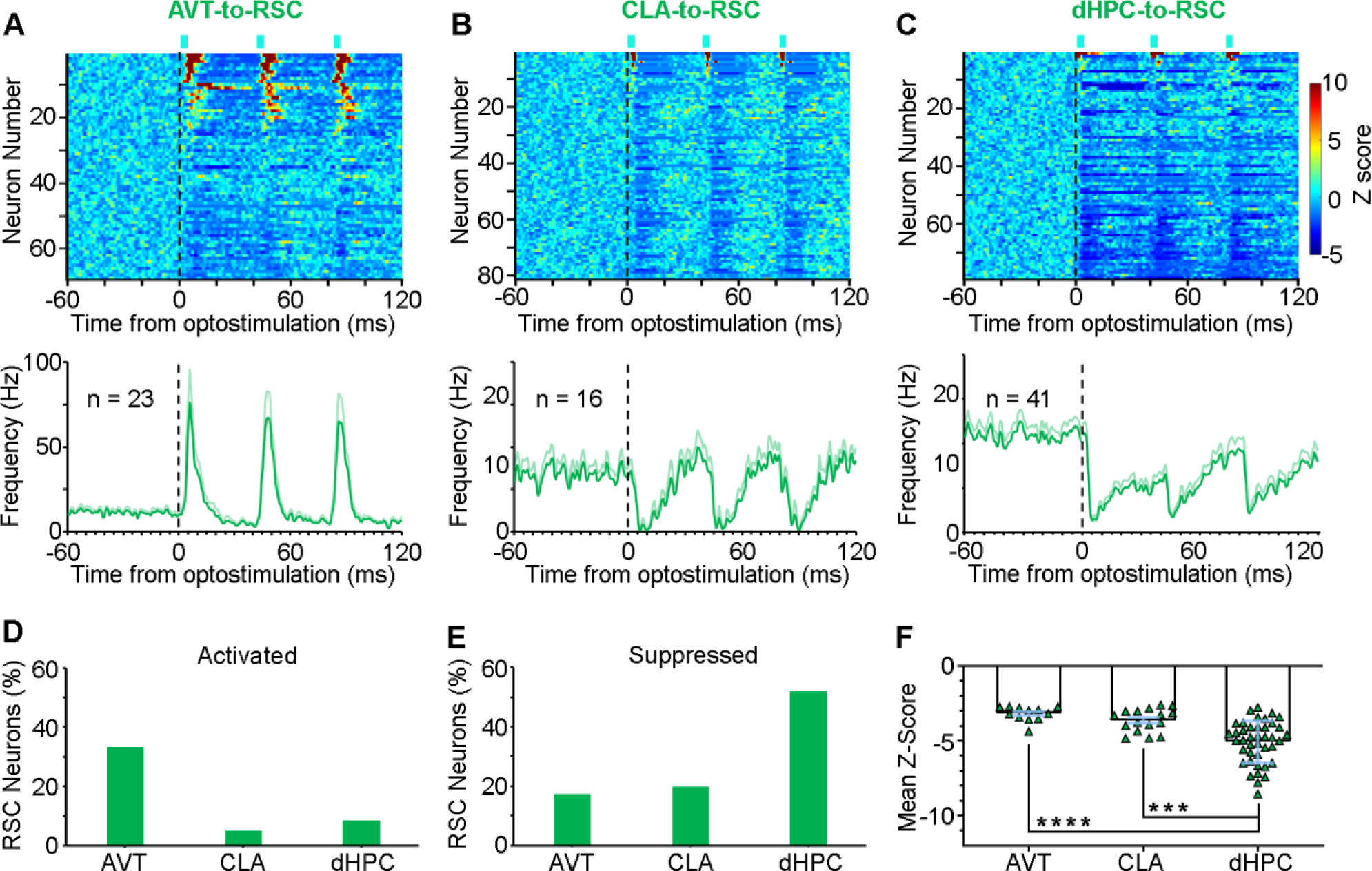
Optostimulation of RSC inputs differentially influence RSC excitatory neurons. (A–C) Peri-event histograms of RSC excitatory neurons in response to optostimulations of the AVT-to-RSC pathway (left), CLA-to-RSC pathway (middle), and dHPC-to-RSC pathway (right). Blue bars: three laser pulses at 25 Hz; pulse width, 3 ms. Top panels, activity heatmap of RSC excitatory neurons. Color bars indicate Z-scored frequency; Z-score transform was based on mean and SD calculated between −60 and 0 ms. Bottom panels, mean activity ± SEM of RSC excitatory neurons activated by AVT optostimulation (left) or suppressed by CLA or dHPC optostimulation (middle and left, respectively). (D&E) Percentage of RSC neurons activated (D) or suppressed (E) by optostimulation. (F) dHPC optostimulation caused significantly greater suppression of RSC neurons compared to CLA or AVT optostimulation.

In comparison, very small portions of RSC putative excitatory neurons were activated by optostimulation of the CLA-to-RSC pathway (8.6%, 7/81; response latency 4.1 ± 1.4 ms, mean ± SD; Figure 6B) or dHPC-to-RSC pathway (5.1%, 4/79; response latency 3.1 ± 1.1 ms, mean ± SD; Figure 6C). These results suggest that the AVT inputs innervate more RSC excitatory neurons compared to the CLA and dHPC inputs (Figure 6D).

Additionally, optostimulation of the inputs suppressed RSC excitatory neurons differently (Figure 6E). Small proportions of RSC putative excitatory neurons showed sustained suppression upon AVT optostimulation (17.4%, 12/69) and CLA optostimulation (19.8%, 16/81; Figure 6B, bottom). In contrast, about half of recorded RSC excitatory neurons were suppressed by dHPC optostimulation (51.9%, 41/79; Figure 6C, bottom). In addition, we found that dHPC optostimulation displayed significantly greater suppression of RSC excitatory neurons compared to AVT and CLA optostimulation (F_(2, 66)_ = 16.846, p < 10^-6^, one-way ANOVA; Figure 6F). This suppression is likely mediated by local interneurons, given the reciprocal connections between RSC excitatory neurons and interneurons (Figure 3). Unfortunately, we were unable to separate the RSC excitatory neurons into DSA and non-DSA neurons due to the relatively small number of simultaneously recorded RSC neurons in most mice, which renders the ICA analysis unreliable. However, it appears that optostimulation of the AVT-to-RSC pathway does not selectively activate the DSA, but rather activates all RSC neuronal types (Suppl. figure 8). Together, these results suggest that RSC excitatory neurons are differentially regulated by inputs from the AVT, CLA, and dHPC, but no input is specifically related to the DSA.

### Optostimulation of RSC inputs similarly influence RSC interneurons

Unlike RSC excitatory neurons, our results revealed that RSC putative interneurons are uniformly activated by optostimulation of the three inputs. Specifically, similar proportions of putative interneurons were robustly activated upon optostimulation of the AVT-to-RSC pathway (68.8%, 22/32; Figure 7A), CLA-to-RSC pathway (66.7%, 14/21; Figure 7B), or dHPC-to-RSC pathway (73.9%, 17/23; Figure 7C). Moreover, the response latency was short for activated RSC interneurons (mean ± SD: AVT, 5.0 ± 1.9 ms; CLA, 5.3 ± 1.0 ms; dHPC, 3.9 ± 1.0 ms). Only a small number of RSC interneurons had a longer response latency upon AVT optostimulation (n = 2; 10 and 12 ms) and dHPC optostimulation (n = 3; 9.5–10 ms). This indicates a primary monosynaptic projection from the AVT, CLA, and dHPC to RSC interneurons, except for a few polysynaptic exceptions.

**Figure 7:**
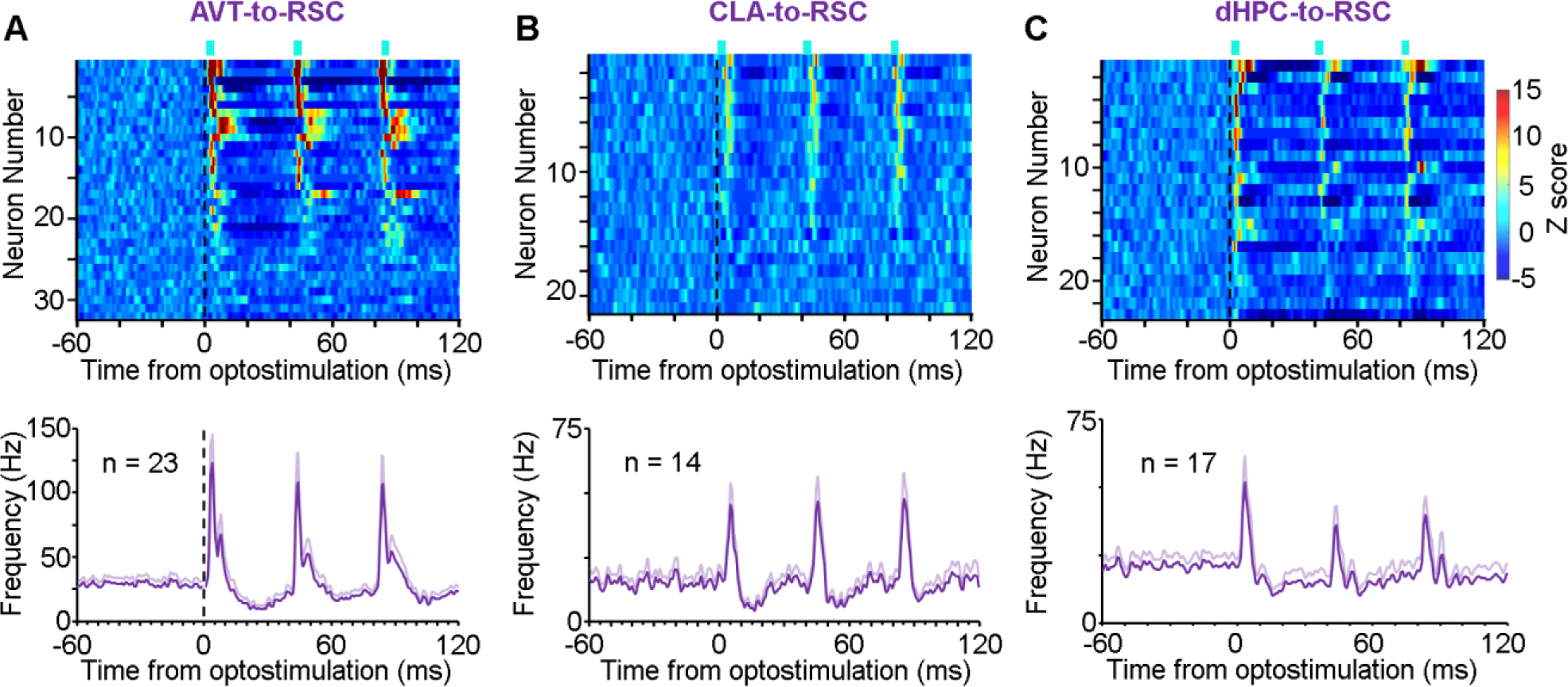
Optostimulation of RSC inputs similarly influence RSC interneurons. (A–C) Peri-event histograms of RSC interneurons in response to AVT (left), CLA (middle), and dHPC (right) optostimulation. Blue bars: three laser pulses at 25 Hz; pulse width, 3 ms. Top panels, activity heatmap of RSC interneurons. Color bars indicate Z-scored frequency; Z-score transform was based on mean and SD calculated between −60 and 0 ms. Bottom panels, mean activity ± SEM of RSC interneurons activated.

## DISCUSSION

We described an assembly of RSC putative excitatory neurons that display relatively high activity during SWS Down states, with maximal firing at the Down-to-Up transitions, which precedes the activity of other RSC neurons. These DSA neurons also display other distinct properties compared to non-DSA excitatory neurons: they have higher firing rates, more burst activity, and likely larger cell body size. Notably, these DSA neurons exhibit higher activity, particularly higher probability of firing during SWS Down states, after a contextual fear learning experience, indicating their role in memory consolidation.

### An exclusive Down-to-Up neuronal assembly

Cortical Down states are thought to represent near-total cortical silence with rare exceptions [11, 36-38]. Here, we report that a considerable portion (∼20%) of RSC deep-layer neurons displays frequent co-activity during the Down state of natural SWS in mice, with maximal activity at the Down-to-Up transition preceding other RSC neuronal activity (Figure 1&2). This RSC DSA is intriguing, as it contrasts with other down state active cortical neurons that display occasional single spike firing during Down states, such as those in the prefrontal, visual, and poster parietal cortices [11, 37, 38].

DSA neurons are positioned to initiate SWS Up states within the RSC for several reasons. Firstly, they fire robustly at Down-to-Up transitions, which precedes the activity of other RSC neurons. Secondly, DSA neurons are localized to the deep layers of RSC. Studies performed *in vivo* and *in vitro* have consistently shown the important role of cortical deep layers, particularly layer 5, in initiating Up states [39-42]. Neurons in layer 5 often fire robustly during Up states and precede neuronal activity in other layers [39, 40]. Furthermore, selective optogenetic activation of layer 5 neurons, but not layer 2/3 neurons, can initiate Up states (Figure 1B&C) and entrain delta oscillations [43], suggesting a causal role of layer 5 neurons in initiating cortical Up states. Thirdly, DSA neurons exhibit significantly higher firing rates and larger spike amplitude compared to non-DSA neurons. It has been shown that higher firing neurons initiate activity at the initiation of Up states [41]. Moreover, larger spike amplitudes could be indicative of larger neurons [27], which have more dendritic surface area for synaptic integration and influence a larger downstream network. These distinct properties of DSA neurons make them suitable for driving SWS Up states.

One potential conceptual framework is that RSC DSA neurons and specific thalamic neurons form a thalamocortical loop to promote cortical Up states [42, 44-46], competing against claustrum-driven cortical Down states [3, 47], thereby generating stochastic Up-and-Down cyclic patterns within the RSC.

### RSC DSA neurons have distinct connectivity

We found that non-DSA (excitatory) neurons and interneurons have reciprocal monosynaptic connections (Figure 3), which is consistent with previous work [18, 29]. However, DSA neurons do not directly communicate with other RSC neurons. Instead, they are likely connected to RSC neurons in different layers and/or extra-RSC brain regions.

A limitation to our study is that the identity of DSA neurons remains unknown. Based on spike waveform and recording sites, DSA neurons are likely layer 5 excitatory neurons, but not interneurons. Therefore, they are likely not ID2+Nkx2.1+ inhibitory neurons, although these interneurons are active during SWS Down states [37]. In addition, DSA neurons are unlikely to be RSC late spiking neurons due to their leading role in the Down-to-Up transition, higher firing activity, and localization to deep layers [48, 49]. It is plausible that DSA neurons are HCN1+ layer 5 neurons, as these neurons exhibited a slow, delta-like rhythm during ketamine-induced dissociation, especially due to their RSC specificity [50].

### RSC DSA and non-DSA neurons support memory consolidation

Previous work has shown that the RSC plays an important role in memory consolidation [14, 19, 23, 24]. Other studies provided initial evidence that manipulating cortical Down states after a new learning experience affects memory consolidation [11, 37]. Here, we directly investigate how distinct RSC excitatory neurons are involved in memory consolidation, particularly during SWS Down-to-Up transitions. We found that RSC DSA neurons exhibit higher firing rates, particularly higher probabilities of firing during post-training SWS Down states. Although this Down-state activity does not necessarily initiate a Down-to-Up transition every time, it likely increases the chances. These findings provide initial evidence supporting the important role of RSC DSA neurons in memory consolidation, likely by promoting SWS Up states.

Additionally, we found that RSC non-DSA neurons increased their activity during memory consolidation. Given that RSC non-DSA neurons and interneurons are often reciprocally connected, they likely interact with each other for information processing, as local interactions between excitatory and inhibitory neurons are considered a hallmark of information coding (24). We speculate that RSC non-DSA neurons facilitate memory consolidation by reactivating specific memory traces, mainly during SWS Up states, but also possibly during Down states when background activity is minimal (Figure 4C) [11]. This is corroborated by recent studies that found RSC memory ensembles displayed increased activity after fear training in mice [51]. Moreover, artificial activation of the RSC tagged ensembles after training induced greater freezing, indicating enhanced memory performance [14]. These effects were only observed when the activation occurred during post-training SWS or anesthesia but not awake [14]. The function of this contextual fear-induced RSC activity increase could be to recruit neuronal ensembles locally in the RSC, along with the network [52].

### RSC neurons are differentially regulated by three primary inputs

An intriguing question remains about what drives DSA activity. Our findings suggest that the AVT, CLA, or dHPC inputs each are unlikely to selectively target the DSA (Figure 5). In reasoning, DSA consists of ∼20% of the RSC neuronal population, while the CLA and dHPC activated < 10% of RSC putative excitatory neurons. Importantly, CLA and dHPC inputs either primarily suppress RSC deep-layer activity or promote SWS Down states [3, 18, 19]. Therefore, CLA and dHPC appear to mainly suppress RSC activity, including that of DSA neurons. Additionally, since the AVT activated ∼33% of RSC putative excitatory neurons, it likely innervates both DSA and non-DSA neurons (Suppl. figure 9), although we cannot exclude the possibility that a specific AVT subpopulation selectively targets RSC DSA neurons [53]. Besides potential innervation by specific inputs, DSA activation may precede other RSC neurons due to intrinsically higher excitability, possibly stemming from distinct membrane properties.

Nonetheless, our data demonstrate that the AVT activates both RSC interneurons and excitatory neurons, while suppressing a small subset of RSC excitatory neurons (Figures 5&6). These findings are supported by several recent studies [17, 48, 54, 55] and indicate that the AVT has a strong influence over RSC neurons. Recent work has shed light on the functional role of the AVT-to-RSC connection, revealing that inhibition of this pathway impairs contextual fear memory formation [17] and memory specificity [54]. These impairments in memory processing may be due to disruption in timing between RSC and hippocampal oscillations relevant in memory formation [55].

Notably, we are the first to report that all AVT^➔RSC^ neurons express Shox2, which thus can be used as a specific marker to target the thalamocortical pathway. As presumed excitatory neurons [33, 35], thalamic Shox2 neurons are positioned to modulate cortical activity through driving rhythms. In support, Shox2 is a transcription factor that has been implicated in rhythmic patterns, not only in thalamus [33], but throughout the body, such as the heart [56] and spinal cord [35].

In contrast, the CLA primarily activates RSC interneurons, which in turn suppress RSC putative excitatory neurons (Figures 5&6). Our findings indicate that the CLA exerts a transient net inhibition of the RSC, which is corroborated by recent study [3]. This CLA-induced inhibition is likely mediated by CLA activation of interneurons [3, 47] or long-range CLA inhibitory neurons [57]. It is important to note that another recent study found that CLA can exert excitatory effects over RSC [58]. This discrepancy could be due to technical differences, regarding the optostimulation protocol (short vs. long) or mouse line (wildtype vs. transgenic). Furthermore, recent work revealed that CLA disruption led to changes in RSC network communication with the prefrontal cortex, along with contextual fear memory processing [59]. This supports the CLA’s ability to orchestrate communication across the cortex due to its vast connectivity [3].

Like the CLA, dHPC primarily activates RSC deep-layer putative interneurons, while suppressing many RSC putative excitatory neurons (Figure 5&6) [18]. This is supported by how hippocampal ripple oscillations – patterns of fast rhythmic neuronal activity – have been shown to preferentially excite RSC excitatory neurons in superficial layer 2/3 but inhibit those in deep layer 5, primarily via the subiculum [18, 19]. Additionally, hippocampal CA1 GABAergic neurons innervate deep RSC excitatory neurons [17], and a small subset of CA1 pyramidal neurons project directly to the RSC [18]. Recent work has focused on the dorsal hippocampus-to-RSC projection, identifying its critical role in memory formation [17, 60, 61] and likely memory consolidation. These findings support the role of dHPC-to-RSC communications in memory consolidation.

Altogether, the AVT, CLA, and dHPC target different sublayers of the RSC [17, 48], supporting the differential effects on RSC neurons, such as proportion of neurons influenced and strength of effects. As an association region, the RSC likely integrates these signals for memory-related processing.

## METHODS

### Mice

A total of 22 mice were used within this study: eight were solely used for *in vivo* recordings, and fourteen were used for optogenetics combined with *in vivo* recordings. Mice were 10–16 weeks at the time of surgery: eighteen of them were C57BL/6J wildtype male mice (Jackson Laboratories, #000664), and four were Shox2::Cre transgenic mice bred in-house [35]). All mice received intra-RSC implantation of 8 or 16 movable tetrodes attached to microdrives (19 and 3 mice, respectively) [18, 62, 63]. Of the 14 mice used for optogenetics combined with *in vivo* recordings, three of them received intra-AVT implantations of optic fibers and intra-RSC microinjections of AAVrg-DIO-ChR2; five of them received intra-CLA implantations of optic fibers and intra-RSC microinjections of AVVrg-ChR2; and six of them received intra-dHPC microinjection of AAV-ChR2 [18].

For comparisons of RSC neurons with other cortical regions, data of ACC neurons were from a previous study [63], and data of PL neurons were recorded from three C57BL/6 male mice. Two additional Shox2-Cre mice were used for characterization of Cre-On/-Off labeling (Suppl. figure 6&7).

After surgery, mice were singly housed in cages (30 x 20 x 20 cm; referred to as home cage) with bedding, a water cup, a cardboard box filled with cotton pads, and wood sticks for environmental enrichment and kept on a 12 h light/dark cycle with ad libitum access to food and water. All procedures were approved by our Institutional Animal Care and Use Committee and were in accordance with the National Research Council’s Guide for the Care and Use of Laboratory Animals.

### Stereotaxic Surgery

Mice were anesthetized with a ketamine/xylazine mixture (∼100/10 mg per kg, i.p.; Vedco Inc.). A bundle or array of 8–16 tetrodes were implanted into the right RSC (AP –1.5 to –1.7 mm, ML +0.4 mm, and DV –0.9 mm). Coordinates for mice with tetrode bundles implanted into the ACC were AP +0.8 mm, ML +0.4 mm, DV – 1.1 mm and PL were AP +1.7 mm, ML +0.3 mm, DV –0.5 mm. For viral injection surgery, wildtype and transgenic mice received injections of AAVrg-CAG-hChR2-H134R-TdTomato (0.4 μL; titer = 1.87 x 10^13 genome copies (gc) per mL; Addgene, #28017-AAVrg) and AAV2rg-EF1a-DIO-ChR2-EYFP (0.4 μL; titer = ∼10^12 gc/mL; NIDA Optogenetics and Transgenic Technology Core), respectively, into the RSC (AP –2.2 mm, ML +0.25 mm, and DV –0.9 mm). Mice that received viral injections underwent a subsequent second surgery two-three weeks later for tetrode array and optic fiber implants. The tetrode arrays were implanted in the RSC with the same tetrode coordinates (above), and an optic fiber (200 μm diameter; ThorLabs, Inc.) was implanted ipsilaterally above either the AVT of transgenic mice (AP –0.45 mm, ML +1.0 mm, DV –2.8 mm at a 10-degree angle) or CLA of wildtype mice (AP +1.1 mm, ML 2.9 mm, DV –2.8 mm). For dHPC optogenetic stimulation combined with recording, see Opalka et al., 2020 [18]. Each tetrode bundle or array was coupled with a microdrive to allow gradual advancement of the electrodes into deeper sites post-surgery [18, 62, 63]. For Cre-On/-Off labeling, transgenic mice received injection of AAV2rg-Ef1a-DO_DIO-TdTomato_EGFP-WPRE-pA (0.2 μL; titer = 8.0 x 10^12 gc/mL; Addgene, #37120-AAVrg) into the RSC and were terminally sacrificed after four weeks.

### Tetrode Recording

Each tetrode consisted of four wires (90% platinum and 10% iridium; 18 μm diameter with an impedance of ∼1–2MΩ for each wire; *California Fine Wire*). About three days after surgery, tetrodes were screened for neural activity. All the analyzed data were recorded at least one week after surgery. Neural signals were preamplified, digitized, and recorded using a Neuralynx Digital Lynx or Blackrock CerePlex acquisition system; the animals’ behaviors were simultaneously recorded. Spikes were digitized at 30 kHz and filtered at 600– 6000 Hz, whereas local field potentials (LFPs) were digitized at 2 kHz and filtered at 1–500 Hz. Ground or a tetrode without clear signal were used as reference for Neuralynx or Blackrock, respectively. All mice received 2–5 sessions of recording when they were freely-behaving or sleeping in home cages (1–4 h). After completion of each recording session, the RSC tetrode array was lowered by 50–100 μm to record a deeper site in the RSC or until DSA detection. We used Plexon Offline Sorter for spike sorting, and sorted spikes were further analyzed in NeuroExplorer (Nex Technologies) and MATLAB (Mathworks). Analyses on RSC spikes were based on a sample size of 175 neurons recorded from eight *in vivo* recording mice, plus two mice also used for optostimulation (28, 21, 18, 18, 17, 16, 15, 14, 14, and 14 neurons simultaneously recorded from each mouse). For optogenetics combined with recordings, analyses on RSC spikes were based on a sample size of 305 RSC neurons recorded from three AVT optostimulation transgenic mice (40, 39, and 22 neurons from each mouse), five CLA optostimulation wildtype mice (46, 18, 18, 10, and 10 neurons from each mouse), and six dHPC optostimulation wildtype mice (34, 26, 19, 18, 3, and 2) [18]. Note that RSC neurons exhibit relatively high frequency firing (5.4 ± 2.3 Hz), which hinders spike sorting; therefore, many not-so-well isolated RSC units were excluded from further analysis. Meanwhile, ACC spikes were based on a sample size of 93 neurons recorded from four mice (33, 30, 16, and 14), and PFC spikes were based on a sample size of 45 neurons recorded from three mice (19, 15, and 11). Low-frequency neurons (< 0.2 Hz) were excluded from analyses due to an insufficient number of spikes.

### Slow-wave sleep and Down state detection

First, detection of rapid-eye movement (REM) sleep and slow-wave sleep (SWS) was determined by the theta (6–10 Hz)/delta (1–4 Hz) ratio extracted from the power spectra of RSC (or ACC/PFC) LFPs when mice remained immobile in their home cage’s ‘bed’ – a cardboard box filled with cotton material, where they exclusively slept. A ratio of 2 or greater was identified as REM, whereas a ratio of 1 or lower was identified as SWS [18, 62, 63]. For Down state detection, RSC (or ACC/ PFC) LFPs were bandpass filtered at 1–4 Hz (MATLAB), and Down (or Up states) were defined as the period when the amplitude of the filtered LFP was > 1 SD above (or < 1 SD below) the mean, when the maximal amplitude is > 2 SD above the mean, and when lasts 0.05–1 s [18, 63].

### RSC assembly detection by combined PCA–ICA

We utilized a previously described combined PCA–ICA method to detect RSC assemblies, subgroups of RSC neurons that exhibited coactivation [26]. Spike counts from natural SWS recordings were binned at 25 ms and z-scored to generate a spike matrix. We then calculated the correlation matrix of the total number of neurons. On the uncorrelated spike matrix, a PCA extracted PCs with significant eigenvalues (coactivations) that surpassed the Marcenko-Pastur distribution threshold, which detected the number of RSC assemblies (Suppl. figure 1B). Next, fastICA computed the independent components (RSC assemblies and its members weights) from the projection of the spike matrix onto the subspace spanned by the PCs with significant eigenvalues. The detected assemblies comprised of a small number of neurons with height weights and a larger number of neurons with small or zero weights. Assembly members were defined as exceeding 1 SD from the mean weight of each assembly. This criterion was used because the normalized weights exceeding 1√N were too generous in assigning assembly members in our recorded RSC population [64]. Meanwhile, 2 SD from the mean weight was too stringent [65], as the same DSA neurons were detected but lesser in number (approximately < 10% of RSC population). The activation strength of each assembly was calculated by projecting the columns of the z-scored spike matrix onto the axis defined by the corresponding RSC assembly pattern. The co-activity events were identified when the activation strength exceeded 2 SD from the mean activation strength of each assembly (Figure 1).

### Spike data analysis

Simultaneously recorded RSC neurons were classified into putative excitatory neuron and interneuron subtypes based on normalized spike waveform between 0–1 ms using PCA analysis [18]. We first extracted the three major principal components (PC1, PC2 and PC3) and then used a hierarchical clustering algorithm (Linkage) to find the similarity (Euclidean distance) between all pairs of spike waveforms in principal components space, iteratively grouping the spike waveforms into clusters based on their similarity. Lastly, we set a distance criterion to extract three clusters from the hierarchical tree (Suppl. figure 2). The burst index for each neuron was defined as the ratio of spikes with peak ISI < 10 ms to the total spikes [53, 66]. Spike amplitude for each neuron was calculated as the difference between the maximum and minimum of the mean waveform. For this analysis, we excluded dendritic waveforms (n = 9) due to the double peak (“W”-shape) confounding the measure [27]. For z-scored firing probability and activity, the Z-score transform was calculated as: Zi = (Xi – x) / s; where Zi and Xi were z-scored and actual values of individual data points, respectively, while x and s were mean and standard deviation (SD), respectively, derived from baseline data points (see legends of Figures 3, 6, & 7).

### Contextual fear conditioning

The fear-conditioning chamber was a custom opaque polycarbonate chamber (25 x 25 x 32cm) with a 36-bar inescapable shock grid floor (Med Associates) [65]. Mice were placed inside the footshock chamber and allowed to explore for 120–180 seconds, at which point a 0.5 s, 0.75 mA foot-shock was delivered. This was repeated four more times at an intertrial interval of 60–180 s. Mice were removed 30 seconds after the last footshock and placed back into their homecage. Note that we used direct-current footshocks to minimize electromagnetic noise and, rarely, mice missed a shock if they stood on two positively or negatively charged grids. The next day, mice were allowed to explore their homecage for ≥ 5 minutes before being re-exposed to the footshock chamber for 5 minutes. Then mice were returned to their homecage for ≥ 5 minutes. (One mouse received the contextual fear recall test 2 hrs after training instead of 24 hours.) Neural activity was recorded during pre-training sleep (∼2 hours), training (∼0.5 hour), post-training sleep (1–2 hours), and contextual fear test (5 min). Half of the mice (n = 3 out of 6) received a cue and trace period prior to the footshock.

### Optogenetic stimulation

Three parameters were used in each session (one, three and five pulses of laser stimulation; 25 Hz; pulse width, 3 ms; laser, ∼1–10 mW, dependent on LFP response to laser) [18]. Pulses of blue laser (473 nm; Opto Engine LLC) were delivered intermittently with an intertrial interval of 10–15 s when mice were freely-behaving or sleeping in home cages (1–2 h). Data from the three and five pulses of stimulation were combined for analyses as shown in Figures 6 & 7. For z-score transform of the correlogram between optostimulation and RSC spikes (bin size = 0.5 ms), mean and SD were calculated between - 60–0 ms: a neuron was considered activated if the post-stimulation peak z-score of at least one pulse exceeded 3.3 (p < 0.001) within 10 ms, while a neuron was considered inhibited if the post-stimulation mean z-score (averaged across the three pulses of optostimulation between 0– 20 ms) was below 2.58 (p < 0.01).

### Histology

Mice received pentobarbital overdose and were deeply anesthetized, and the final electrode position was marked by passing a 10 mA, 20 s current through two or each tetrode. Then, mice were perfused transcardially with 1× PBS followed by 10% formalin. Brains were removed and post-fixed in 10% formalin, allowing for ≥ 24 h before vibratome slicing (50 mm coronal sections; Leica VT1000 S). Coronal sections were mounted with a Mowiol mounting medium (Mowiol 4-88, Calbiochem) based on a recipe from Cold Spring Harbor Protocols (pdb.rec10255) and imaged for verification of all recording sites, injection sites, viral expression, and optical fiber placements. Representative images in Figure 5 were acquired with Leica DM5500B fluorescent microscope, Suppl. figure 7 were acquired using Leica Stellaris confocal microscope during a demonstration, and Suppl. figure 8 was acquired with Leica DM6 Thunder fluorescent microscope using THUNDER Imager computational clearing.

### Statistics

Sample sizes were based on similar studies from our lab [18, 67]. All statistical analysis was calculated in SPSS. Unless otherwise stated, non-parametric Wilcoxon rank sum, Wilcoxon signed-rank, and Kruskal–Wallis one-way analysis of variance (ANOVA) on ranks tests were used. When parametric tests were used, the assumptions for normality and homogeneity of variances were met (Shapiro-Wilk and Levene’s tests, respectively).

## SUPPLEMENTAL FIGURES

**Suppl figure 1:**
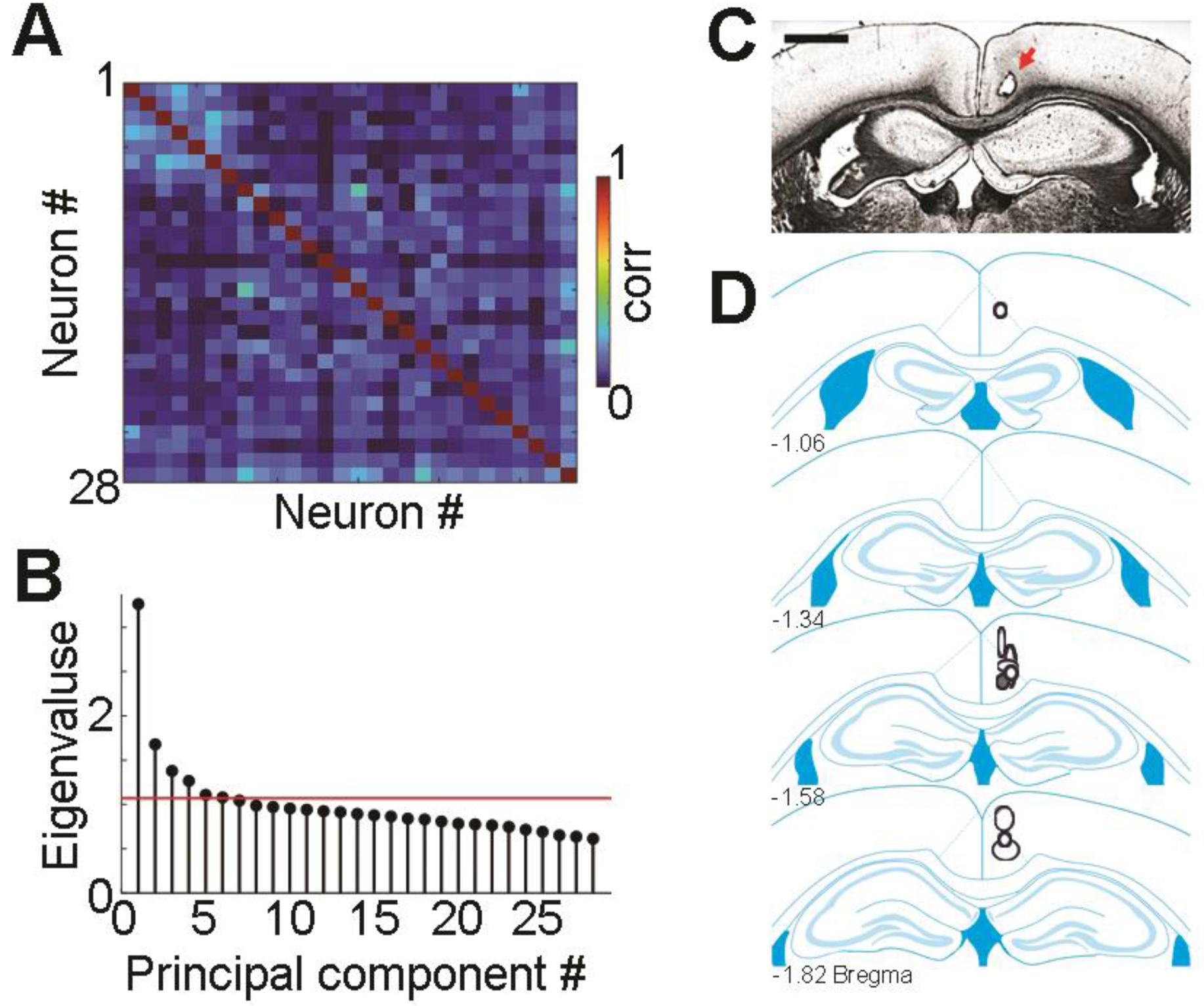
DSA is localized towards the granular RSC. (A&B) Spike train correlation matrix (A) and detection of assembly numbers using Marchenko– Pastur threshold (B) from example in Figure 1A&B with neurons arranged in the same order. (C&D) Example coronal section (C) and schematics of final recording sites (D). Note larger shapes indicate multiple tetrode lesion sites. Shaded circle represents recording site from example in C. Scale bar = 1 mm.

**Suppl figure 2:**
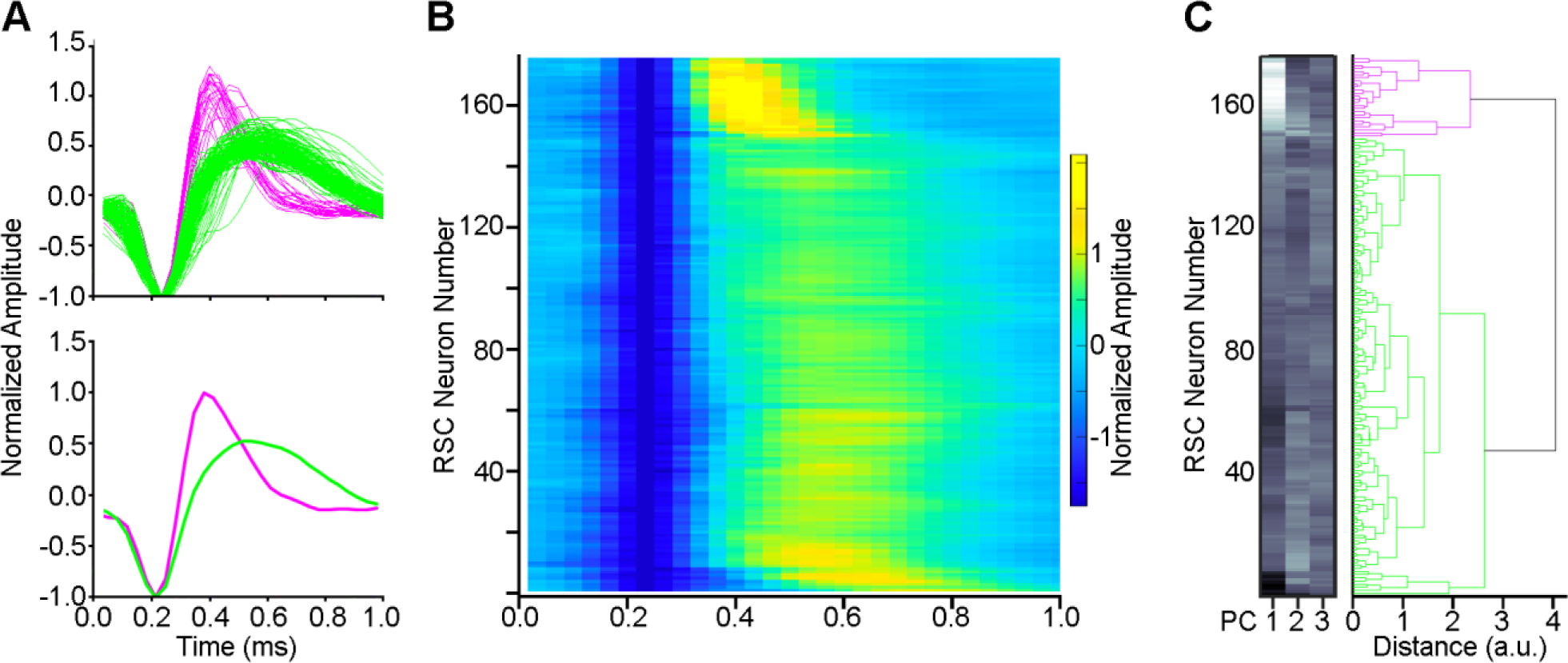
RSC putative inhibitory and excitatory neurons classified based on spike waveform using principal component analysis (PCA). (A) Normalized amplitude of RSC neurons individually (top; n = 175) and averaged (bottom). (B) Heatmap of spike waveforms from the same RSC neurons. (C) PCA classified RSC neurons based on the first three principal components (PCs), coded from low scores (white) to high scores (black). Our PCA classified RSC putative interneurons (magenta, n = 26) exhibiting narrow spike waveforms. RSC putative excitatory neurons (green, n = 149) exhibited wider spike waveforms compared to interneurons.

**Suppl. figure 3:**
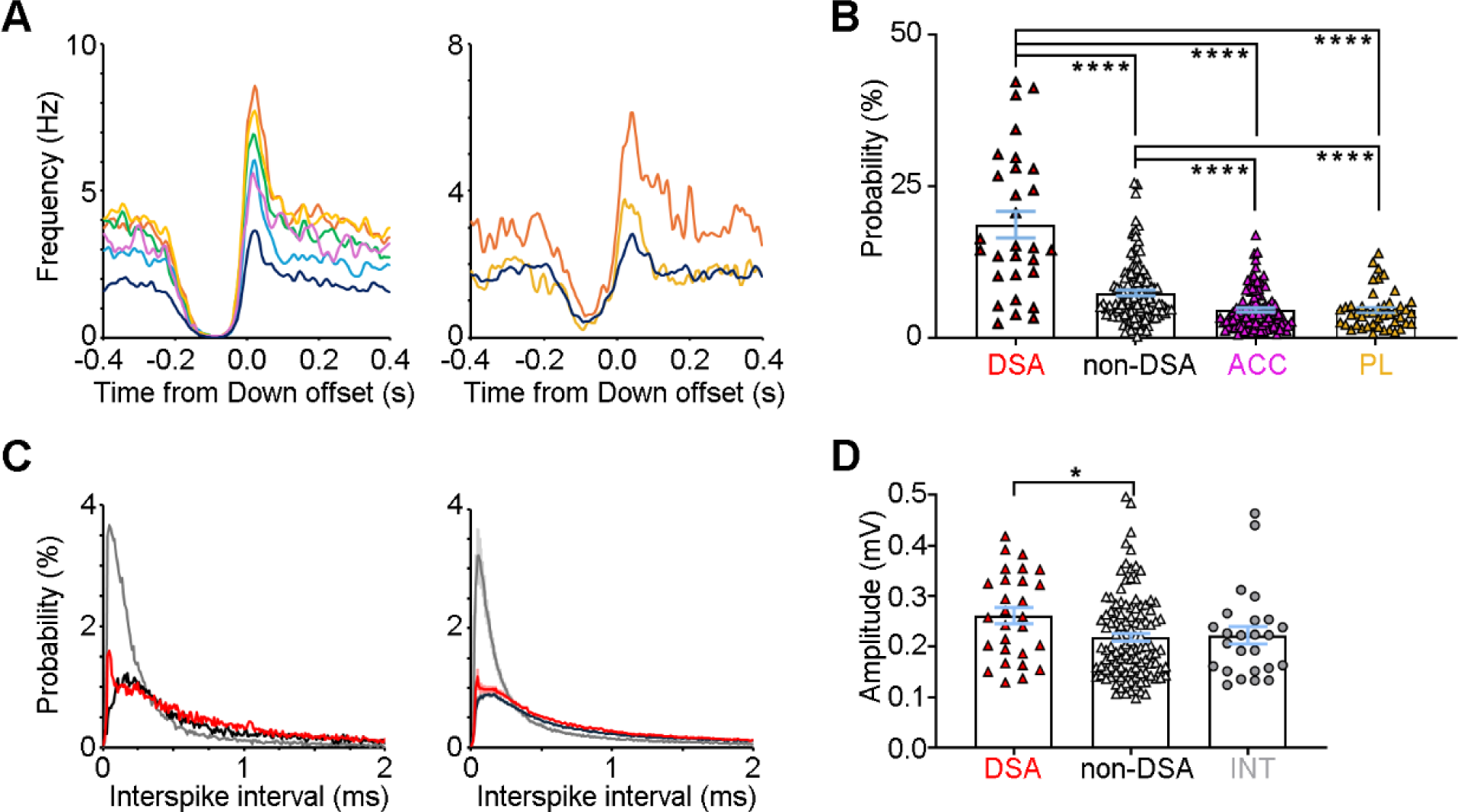
DSA neurons are distinct from other cortical neurons. (A) Mean frequency of ICA-detected neuronal assemblies (colors) and other neurons (navy) from an example mouse in the ACC (left) and PL (right). Note that no assembly activity precedes other neuronal activity. (B) RSC DSA neurons exhibit significantly greater down state firing probability compared to RSC non-DSA neurons, ACC neurons, and PL neurons (*H*_(2)_ = 67.157, p < 10^-14^, KW test). Note ACC and PL each only had one interneuron, which was included. (C) Inter-spike interval of DSA neurons (red), non-DSA neurons (black), and interneurons (left, examples) and across mice (right, n =10, mean ± SEM). (D) DSA neurons have significantly greater spike amplitude compared to non-DSA neurons but not interneurons. Error bars = mean ± SEM.

**Suppl figure 4:**
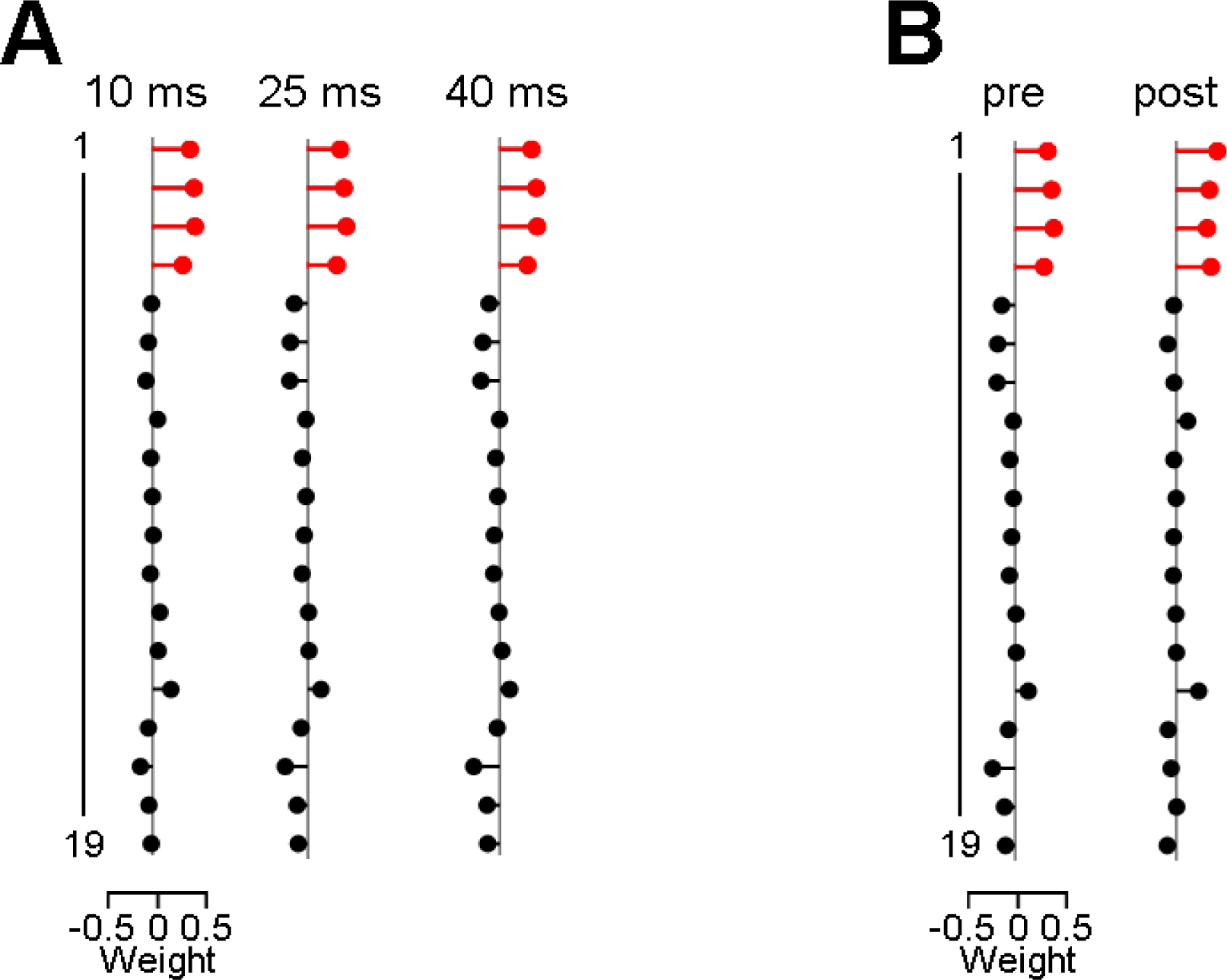
DSA membership is robust, using different bin sizes and during contextual fear conditioning paradigm. (A) An example independent component analysis (ICA) detected the DSA (red) at varying bin sizes of 10 ms (left), 25 ms (middle), and 40 ms (right). Note that in a few situations, changing the bin size led to inclusion or removal of 1–2 DSA neurons (3 mice) or in/decrease of the number of assemblies detected by 1 (3 mice). (B) DSA membership remains consistent across pre- and post-training SWS (25 ms bins, pre is the same as A, middle).

**Suppl. figure 5:**
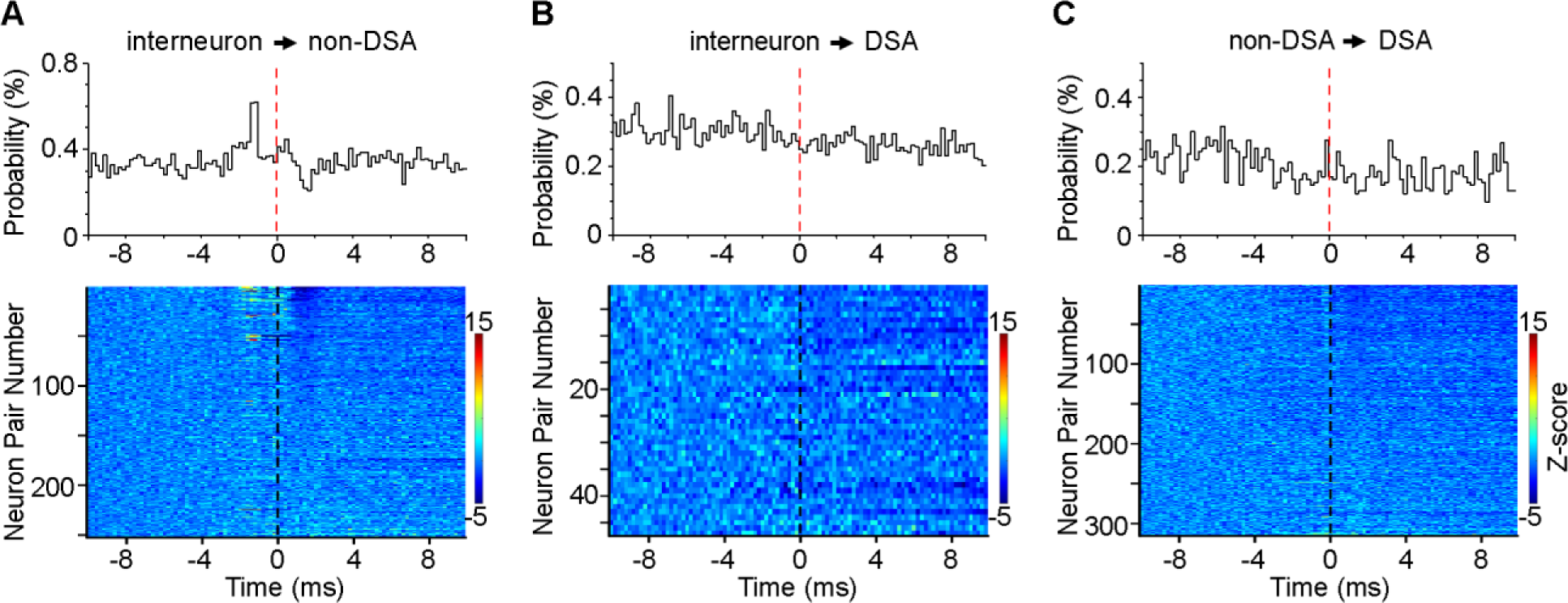
DSA neurons receive weak local connections. (A–C) Cross-correlations between RSC neuronal subpopulations. Neuron pairs include interneurons and non-DSA neurons (A), interneurons and DSA neurons (B), and non-DSA and DSA neurons (C). non-DSA neurons were used as the reference in A, while DSA neurons were the reference in B and C. Top panels, representative neuronal pairs. Bottom panels, heatmap of individual neuron pairs. Color bar indicates Z-scored firing probability. Neurons are arranged from high-to-low firing between 1–2 ms, and Z-score transform was calculated between -10 and -5 ms based on mean and SD.

**Suppl. figure 6:**
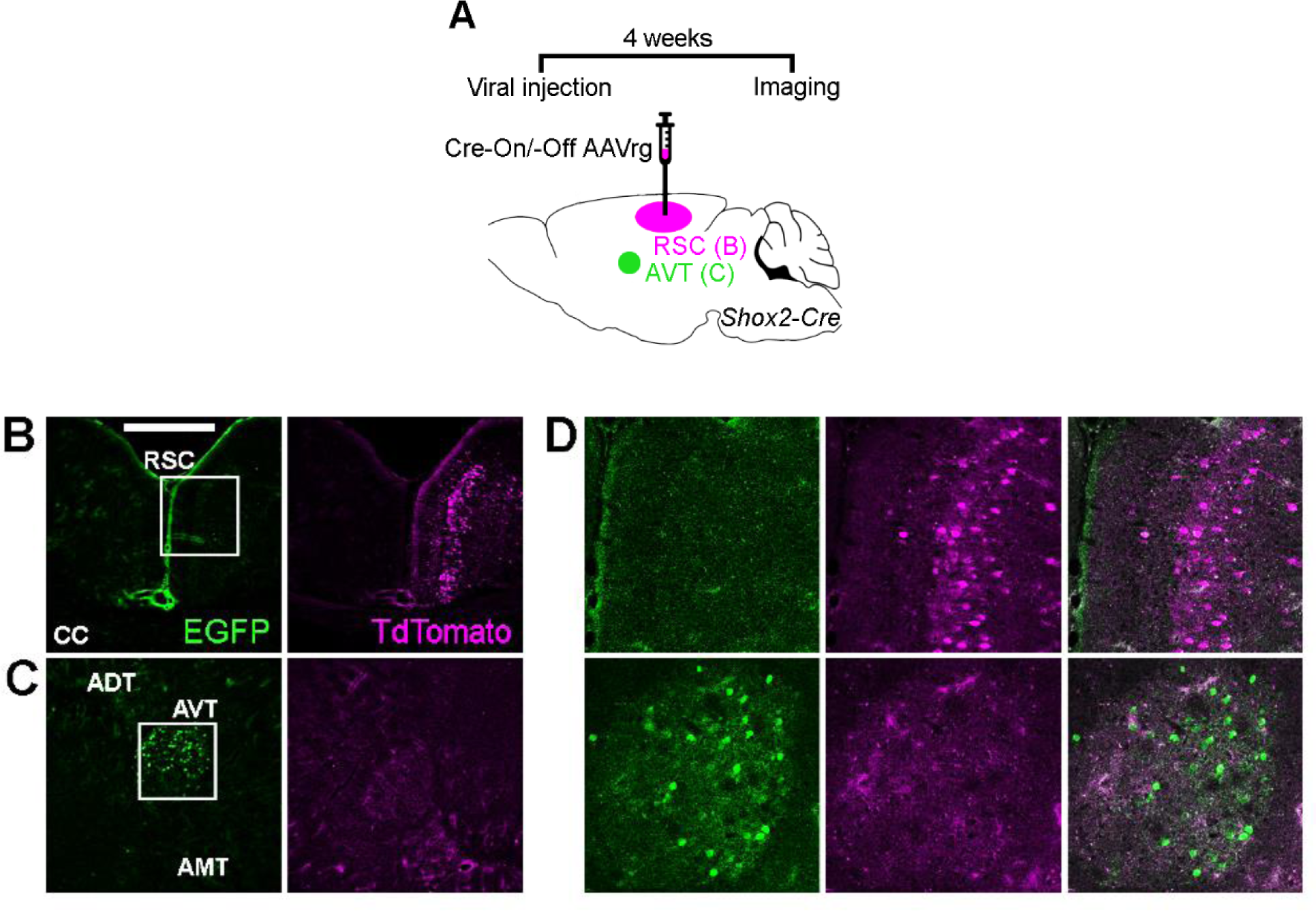
An approach to selectively target the AVT-to-RSC pathway. (A) Top, surgical timeline. Bottom, schematic drawing of AAV-retrograde (rg)-Cre-On/-Off virus (0.2 μL) injected in granular RSC and retrograde labeling in AVT of adult Shox2-Cre mice, which enables the expression of EGFP (or TdTomato) when Cre is present (or absent). (B&C) Representative coronal sections (from the same brain) showing EGFP+ terminal labelling (B, left) and TdTomato+ cell bodies (B, right) in RSC injection site and EGFP+ (C, left) but no TdTomato+ cell bodies (C, right) in AVT. Scale bar, 500 μm (D) Top, insets from B showing AAV-retrograde (rg)-Cre-On/-Off viral expression in granular RSC. Bottom, inset from C showing retrograde labeling of EGFP reporter protein in AVT. CC, corpus collosum; ADT, anterodorsal thalamus; AMT, anteromedial thalamus.

**Suppl. figure 7:**
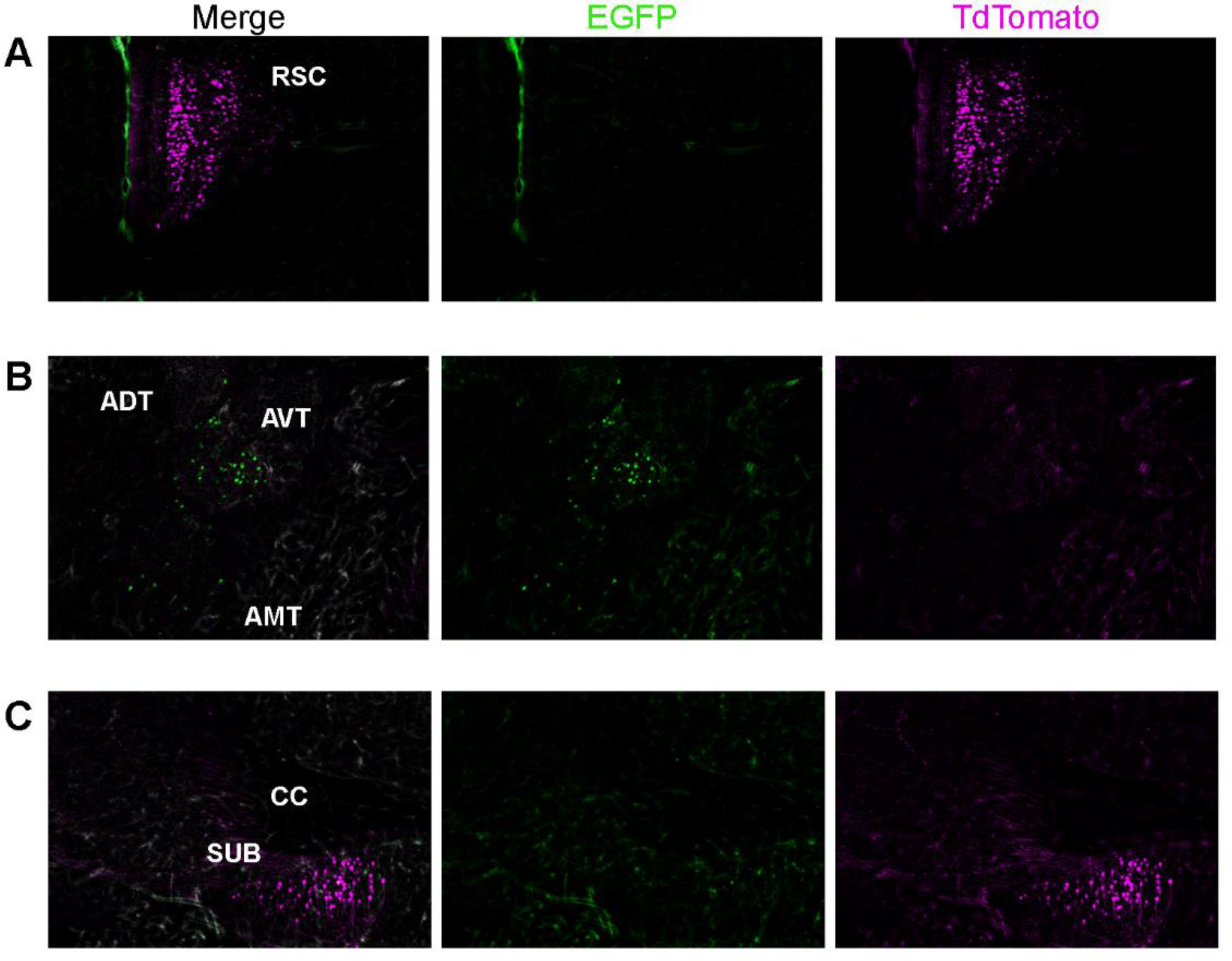
AVT^➔RSC^ but not SUB^➔RSC^ neurons express the transcription factor Shox2. (A–C) Representative coronal sections (from the same brain) of AAV-retrograde (rg)-Cre-On/-Off virus (0.2 μL) injected in granular RSC (A) and retrograde labeling in AVT (B) and subiculum (SUB; C) of adult Shox2-Cre mice, which enables the expression of EGFP (or TdTomato) when Cre is present (or absent). RSC injection site (A) displays EGFP+ terminal labelling and TdTomato+ cell bodies. Meanwhile, AVT (B) displays EGFP+ but no TdTomato+ cell bodies, whereas SUB (C) displays TdTomato+ but no EGFP+ cell bodies. ADT, anterodorsal thalamus; AMT, anteromedial thalamus; CC, corpus collosum.

**Suppl. figure 8:**
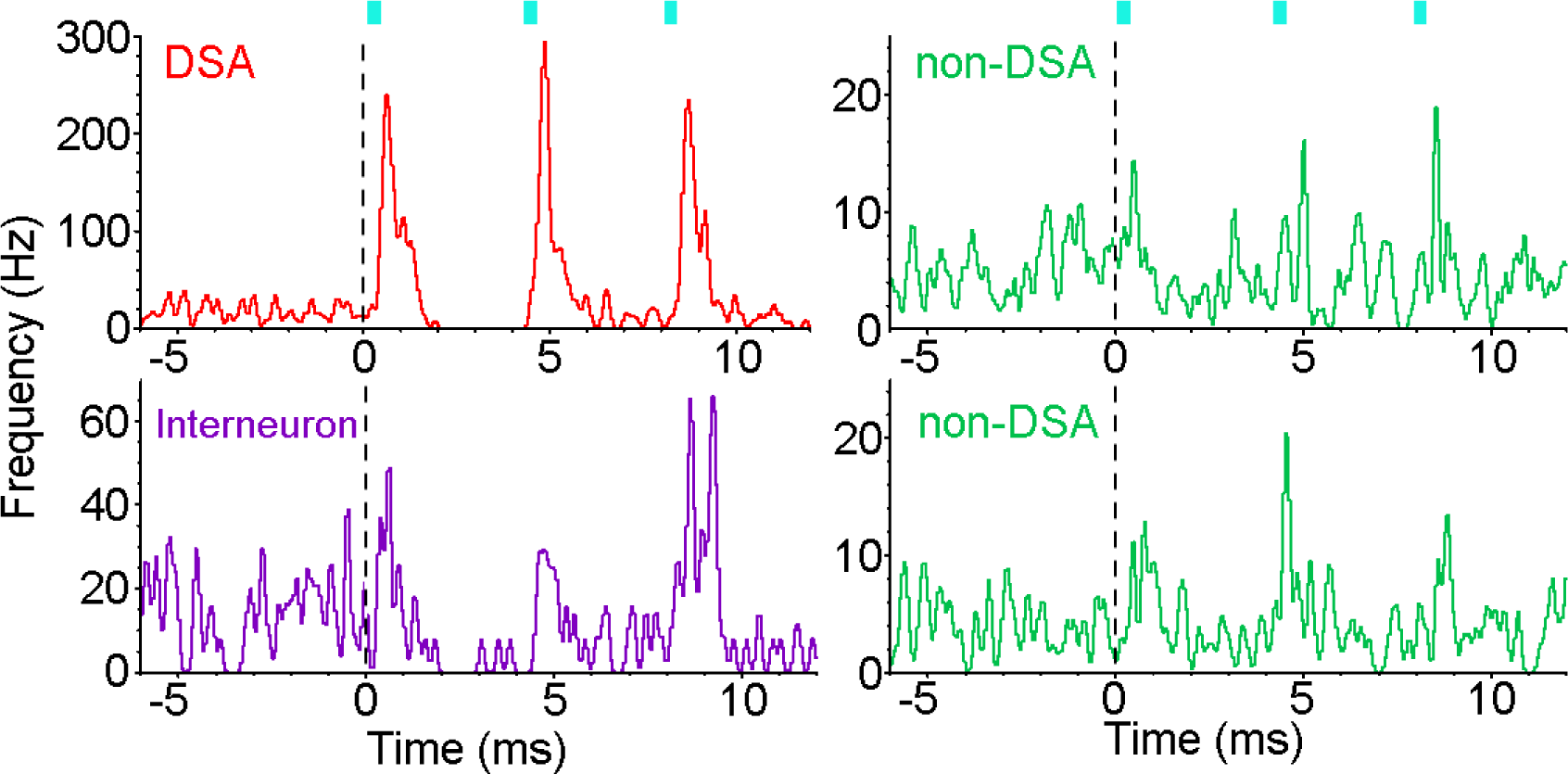
AVT-to-RSC optostimulation activates each RSC neuronal subpopulation. Peri-event histograms of four RSC neurons activated by AVT optostimulation. Blue bars: three laser pulses at 25 Hz; pulse width, 3 ms. Note that AVT-to-RSC optostimulation strongly activated a DSA neuron (red) but also an interneuron (purple) and non-DSA neurons (green) to a lesser extent.

